# The N-terminal Segment of the Human ANKZF1 negatively regulates its internal mitochondrial targeting signal to prevent its mitochondrial localization

**DOI:** 10.1101/2025.04.04.647180

**Authors:** Mudassar Ali, Bhoopesh Maheswaran, Devid Sahu, Nidhi Malhotra, Koyeli Mapa

## Abstract

ANKZF1, the human orthologue of yeast Vms1, is a multi-domain cytosolic protein. Occasionally, the protein is also found in mitochondria, although the reason for its mitochondrial localization and the mechanism of its mitochondrial targeting remains unclear. Despite the absence of any predicted Mitochondrial Targeting Sequence or Domain (MTS/MTD) in the protein, ANKZF1 possesses multiple internal Matrix-targeting sequence-like sequences (iMTS-Ls). In this study, we show that ANKZF1’s N-terminal 73 residues negatively regulate its mitochondrial targeting. Interestingly, 1-72-ANKZF1 peptide hinders the mitochondrial targeting of Δ73-ANKZF1 upon co-expression. Using a series of truncation mutants of ANKZF1, we further show that iMTS-Ls constituted by residues 231-240 of ANKZF1 is indispensable for its mitochondrial localization. The sequence of 231-240 residues is extremely conserved across different organisms, indicating the importance of this segment for conditional mitochondrial targeting of the protein. Importantly, 231-324 residues containing two consecutive predicted iMTS-Ls constitute an independent mitochondrial signal sequence and can target Green Fluorescent Protein (GFP) to mitochondria when fused to N-terminus. Further, by Molecular Dynamic simulation, we show that the deletion of N-terminal 74 amino acids of ANKZF1 leads to a massive structural rearrangement within the protein leading to the opening of its C-terminal part and solvent exposure of the 231-240 residues. We posit that these structural rearrangements and exposure of the internal mitochondrial targeting sequence in the absence of N-terminal segment of ANKZF1 leads to its mitochondrial translocation.

## Introduction

Mitochondria are essential organelles of eukaryotic cells which serve as the hub of key metabolic pathways. All enzymes of these metabolic pathways and most of the mitochondrial constituent proteins except a few subunits of the respiratory chain complexes are encoded by the nuclear genome despite the presence of mitochondrial DNA. The nuclear genome-encoded mitochondrial precursor proteins, also known as preproteins, are synthesized in the cytoplasm and are subsequently translocated to mitochondria. For translocation across the mitochondrial membranes and intra-mitochondrial sorting of the preproteins, there are elaborate machinery present on both the mitochondrial membranes. Different translocases, popularly known as TOM (Translocase of Outer Mitochondrial membrane) and TIM (Translocase of Inner Mitochondrial membrane) complexes, collaborate to send the passenger proteins to their final intra-mitochondrial destination. For about half of the resident proteins of mitochondria, the nature of Mitochondrial Targeting Signals (MTS) is known. Over the last years, various computational prediction tools like TargetP (1), mitoFates (2), TPred3 (3), and DeepMito (4) have been developed using deep learning, to identify potential MTSs within any protein. For most of the matrix-targeted proteins, there is a well-defined N-terminal pre-sequence, also known as Mitochondria-Targeting Signals or Sequences (MTS). The pre-sequences most commonly constitute amphipathic helices. For a certain percentage of mitochondrial resident proteins, the targeting signals are not well defined and are difficult to predict by in-sillico prediction tools. Recently, many cellular proteins have been described with dual or multiple cellular localization which localize to mitochondria under specific conditions, especially during various stresses. Understanding conditional mitochondrial targeting of various proteins has elucidated a plethora of fascinating mechanisms employed by different cellular proteins for alternate sub-cellular localizations to perform specific biological functions.

Human ANKZF1 is the yeast orthologue of Vms1 (5,6). Recent literature has shown the important roles of Vms1/ANKZF1 in the Ribosome Quality Control (RQC) pathway as tRNA hydrolase and releases nascent polypeptide chains from stalled ribosomes (7,8). Importantly, ANKZF1 overexpression is associated with the progression of many cancers (9–12), although the role of the protein in cancer pathogenesis or disease progression, has not been explored so far. ANKZF1 is a multidomain protein of 726 amino acids. WT ANKZF1 has an N-terminal part with many Leucine residues, and in its yeast orthologue, Vms1, the equivalent segment is a Leucine-Rich Sequence (LRS). However, there are many more leucine residues throughout the ANKZF1 sequence. The N-terminal segment is followed by a Zinc-finger domain (ZnF). Next to the ZnF domain, there is a putative MTD (Mitochondrial Targeting Domain), the exact length of which is not known. This domain also contains the VLRF (Vms1-like Release Factor) domain. The putative MTD/VLRF is followed by two ankyrin repeats and a coiled-coil motif. In a recent study, we have shown that ANKZF1 mostly remains in the cytoplasm except for certain mitochondrial stresses (13). ANKZF1 is recruited to damaged mitochondria during proteotoxic stress due to the expression of misfolded or aggregated proteins within mitochondria or due to depolarization of mitochondria, specifically in the presence of Parkin (13).

Occasionally, we have also observed ANKZF1 to be co-localized with the mitochondrial network in the absence of any given stress, although the reason for its mitochondrial localization remains elusive so far. Previous literature has shown the localization of ANKZF1 to mitochondria during oxidative stress like its yeast Orthologue, Vms1 (14). Vms1’s N-terminal LRS sequence was shown to interact with its MTD and prevent its mitochondrial localization (15).

In the current study, we show that ANKZF1 contains an atypical internal mitochondrial signal sequence that remains hidden due to intra-molecular interaction with the N-terminal part of the protein. Truncation of N-terminal 73 amino acids of ANKZF1 leads to its mitochondrial localization, indicating that 1-73 amino acids negatively regulate the mitochondrial localization of the protein. Interestingly, 1-72-ANKZF1 peptide efficiently and specifically prevents the mitochondrial targeting of Δ73-ANKZF1. Furthermore, serial truncations of ANKZF1 show that residues 231-240 are indispensable for its mitochondrial targeting, and this segment is evolutionarily conserved. Moreover, we delineated that 231-324 amino acids can act as an independent Mitochondrial-Targeting Sequence (MTS), and a Green Fluorescent Protein (GFP) fused to these 94 amino acids at the N-terminus is efficiently targeted to mitochondria. To further understand the contribution of the N-terminal part of ANKZF1 on its structural features and ultimately on mitochondrial translocation, MD simulation of the wild-type (WT) protein and the N-terminal 74 residue-truncation mutant was carried out. Notably, the WT protein shows the interaction of the VLRF domain (203-346 residues) containing the MTS with the N-terminal part (1-74 residues) as well as the C-terminal part (600-726) of the protein. Interestingly, Δ74-ANKZF1 mutant shows a massive structural reorganization compared to the WT protein. In the truncation mutant, the C-terminal part completely changes its orientation and loses its interaction with the VLRF domain. This structural reorganization leads to solvent exposure of 231-240 residues, the indispensable part of ANKZF1’s internal MTS, leading to the mitochondrial localization of the protein.

## Results

### ANKZF1 possesses a cryptic mitochondrial targeting signal

To understand the molecular basis of ANKZF1’s occasional mitochondrial localization (Figure S1A), we made two N-terminal truncations; the first truncation is without the N-terminal 73 amino acids, and the second truncation is devoid of N-terminal 210 amino acids. We reasoned that a similar interaction of the N-terminal part of ANKZF1 on its putative MTD known for its yeast ortholog, Vms1, may prevent its mitochondrial localization (16). Δ210-ANKZF1 was made, keeping in mind the fact that the second other major variant of ANKZF1 that is found apart from the full-length protein starts from 211 residues. The full-length and N-terminally truncated ANKZF1 proteins (Δ73 and Δ210) were C-terminally fused to GFP to assess the sub-cellular localization of these truncated proteins (Figure 1A). HeLa cells were transfected with the full-length and truncated ANKZF1 constructs, and 24 hours after transfection, cells were treated with MitoTracker-Red to stain the mitochondria, and images were captured by confocal microscope. Wildtype (WT) ANKZF1 protein shows diffused cytosolic distribution; however, both Δ73-ANKZF1 and Δ210-ANKZF1 proteins were found to localize to mitochondria (Figure 1B and 1C). As both truncations showed complete overlapping signals with MitoTracker, it suggested the presence of a cryptic mitochondrial targeting domain/signal in ANKZF1.

**Figure 1:**
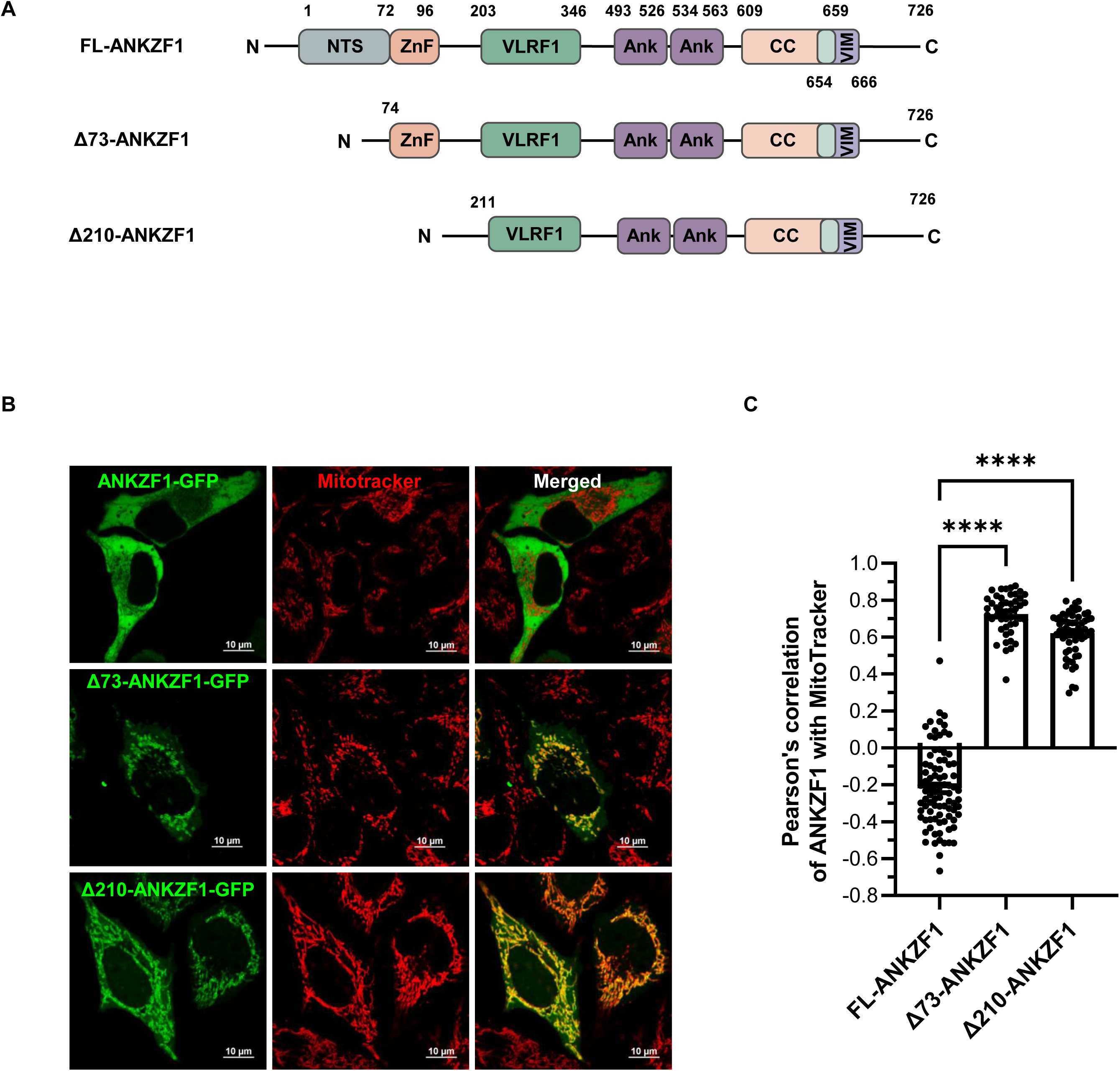
ANKZF1 possesses a cryptic mitochondrial targeting domain. **A.** Schematic overview of ANKZF1 domain map of full length WT protein consisting of N-terminal segment (NTS), ZnF (Zinc finger domain), VLRF domain, two ankyrin repeats followed by coiled-coil motif, and VCP interacting motif. The N-terminally truncated variant lacks the initial 73 amino acids termed as Δ73-ANKZF1 and another variant lacks the NTS and ZnF and is termed as Δ210-ANKZF1. The numbers represent the residue numbers for each domain. **B.** Cellular localization of full-length ANKZF1, Δ73-ANKZF1 and Δ210-ANKZF1 C-terminally tagged with EGFP in HeLa cells was checked by confocal microscopy. Mitochondria were stained with MitoTracker deep red. WT ANKZF1 shows cytosolic localization while both the truncated variants show mitochondrial localization. **C.** Pearson’s correlation coefficient of co-localization of ANKZF1-GFP (FL-ANKZF1-GFP, Δ73-ANKZF1-GFP and Δ210-ANKZF1-GFP) and MitoTracker deep red was calculated. Values represent means ± SEM, N=3, Kruskal-Wallis test with Donn’s multiple comparisons test was performed to determine the mean differences, **** indicates P < 0.0001.

### Amino acid residues 231-240 of ANKZF1 are essential for its mitochondrial localization

Mitochondrial localization of Δ210-ANKZF1 indicated that the MTD of ANKZF1 must lie after 210 amino acids from the N-terminal. To assess the position of MTD, we made four constructs with N-terminal truncations; Δ250-ANKZF1, Δ330-ANKZF1, Δ370-ANKZF1, and Δ410-ANKZF1 were generated with a GFP fusion tag at C-terminus of the protein (Figure S1B). All the truncation mutants were transfected in HeLa cells and cellular localization was checked by imaging by confocal microscopy following MitoTracker staining. None of the truncation mutants (Δ250-ANKZF1, Δ330-ANKZF1, Δ370-ANKZF1, and Δ410-ANKZF1) showed mitochondrial localization (Figure S1B, Figure S1C and S1D). Δ210-ANKZF1 was used as the positive control for mitochondrial localization (Figure S1C, uppermost row). This data indicated that the MTD of ANKZF1 lies within 210 and 250 residues. Next, to precisely determine the MTD, we narrowed down the N-terminal truncations further by ten amino acids and generated Δ220-ANKZF1, Δ230-ANKZF1, and Δ240-ANKZF1 with GFP fusion tag at the C-terminal (Figure 2A). Interestingly, Δ220-ANKZF1 and Δ230-ANKZF1 truncation mutants showed mitochondrial localization (Figure 2B, 2^nd^ and 3^rd^ rows from top and Figure 2C), while Δ240-ANKZF1 showed diffused cytosolic localization (Figure 2B, 4th row from top and 2C). Here, Δ210-ANKZF1 and Δ250-ANKZF1 were used as positive and negative controls for mitochondrial localization, respectively (Figure 2B, top and bottom rows, respectively). This result confirms the presence of MTD of ANKZF1 between 231 to 240 residues (Figure 2B). ANKZF1 sequence analysis shows this 10-residue stretch is a highly conserved region among the vertebrates and in *Saccharomyces cerevisiae* (Figure 2D). The synthesis of N-terminal truncated proteins was confirmed by western blot using the anti-GFP antibody, showing the expression of truncated variants at the expected molecular weights along with the full-length protein (Figure S2A).

**Figure 2:**
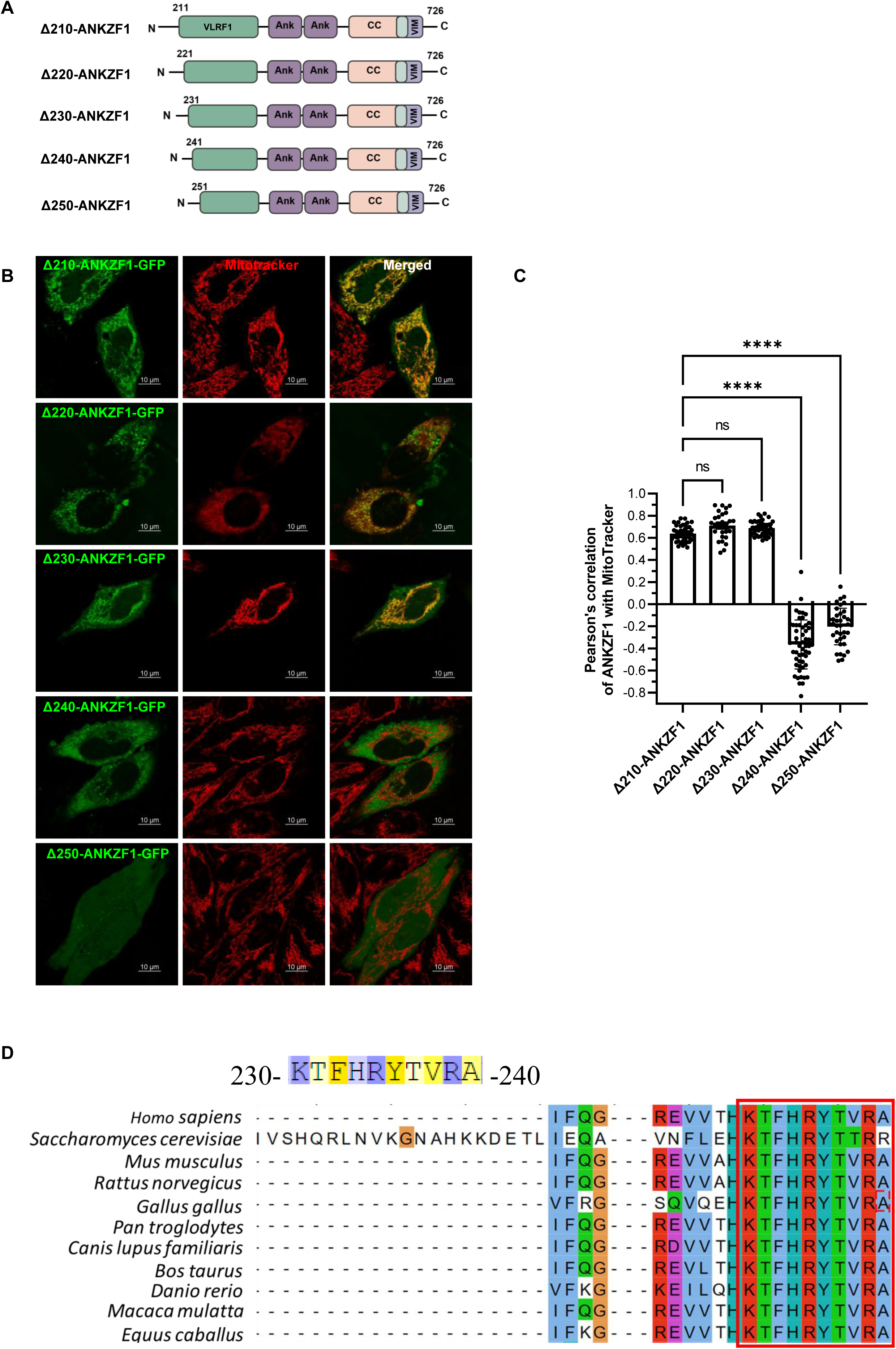
ANKZF1 possesses a non-canonical mitochondrial targeting signal. **A.** Schematic representation of ANKZF1 truncation mutants from 210 to 250 amino acids with each 10 amino acid span, Δ210-ANKZF1, Δ220-ANKZF1, Δ230-ANKZF1, Δ240-ANKZF1, and Δ250-ANKZF1. **B.** Confocal microscopy images show the localization of GFP tagged ANKZF1 truncation mutants. Δ210-ANKZF1-GFP is used as positive control for mitochondrial localization while Δ250-ANKZF1-GFP construct was used as a negative control for mitochondrial localization. Δ220-ANKZF1 and Δ230-ANKZF1 truncated variants show mitochondrial localization and co-localized with MitoTracker deep red. However, the Δ240-ANKZF1 variant shows cytosolic localization. **C.** Bar graph represents Pearson’s correlation coefficient of co-localization of ANKZF1-GFP (Δ210-ANKZF1, Δ220-ANKZF1, Δ230-ANKZF1, Δ240-ANKZF1, and Δ25ANKZF1) and MitoTracker deep red. Values represent means ± SEM, N=3, one-way ANOVA with Bonferroni’s multiple comparisons test was performed to determine the mean differences, **** indicates P < 0.0001. **D.** Peptide sequence from 230-240 amino acids of ANKZF1 is shown at top. Multiple sequence alignment of amino acids 230-240 of different organisms shows the evolutionary conservation of this region of the protein throughout the eukarya.

### ANKZF1 harbors multiple predicted Internal MTS-like (iMTS-L) sequences and two consecutive iMTS-Ls at amino acids 231-324, constitute an independent mitochondrial targeting sequence

WT ANKZF1 did not show any predicted MTS by commonly used prediction algorithms like TargetP (1), mitoFates (2), TPred3 (3) or DeepMito (4). As the N-terminal truncations showed mitochondrial localization, we also checked the predicted localization of truncation mutants using the same tools. Although mitoFates, TPred3 or DeepMito gave similar results for truncation mutants, MULocDeep (17,18) revealed ∼70% propensity for mitochondrial localization of the Δ220-ANKZF1 (Figure S2B). However, the WT protein did not show any prominent mitochondrial localization with MULocDeep. Recent literature showed that many proteins that are found in mitochondria do not harbour conventional mitochondrial targeting signals. Instead, these proteins contain internal Matrix Targeting Signal-like sequences (iMTS-Ls) (19–21). To check the presence of iMTS-Ls in the ANKZF1 sequence, we used iMLP tool described in Schneider et al (21) and found 6 prominent iMTS-Ls throughout the primary sequence of ANKZF1 (Figure 3A). Interestingly, amino acids 200-250 and 270-325 showed two predicated high propensity iMTS-Ls (Figure 3B). As shown in the previous section, 231-240 residues present in the first high-propensity iMTS-Ls, are well conserved across different species and are indispensable for mitochondrial localization. Although 270-325 residues are predicted to constitute another high propensity iMTS-Ls, we have shown that these iMTS-Ls or the ones at more at the C-terminal part, are not sufficient to translocate the protein to mitochondria. To check whether these predicted iMTS-Ls can act as independent mitochondrial targeting signals (MTS), we fused the residues 221-250, 231-282 and 231-324 of ANKZF1 in the N-terminus of GFP. Among these three segments, only 231-324-ANKZF1 was able to target GFP to mitochondria (Figure 3C). 221-250-ANKZF1 and 231-282-ANKZF1 although they contain the indispensable residues (231-240) for mitochondrial translocation, these two peptides were unable to act as an independent targeting signal (Figure 3C). Using alpha-fold, we checked the predicted structure of the peptide 231-324-ANKZF1 that constitutes a complete MTS. 231-324-ANKZF1 adopts majorly an alpha-helical structure with a preceding beta-sheet in the N-terminus and two beta-sheets at the C-terminal part of the peptide (Figure 3D). The Matrix Targeting pre-sequences or the iMTS-Ls are known to form alpha helices although apart from alpha helix, we observed three short beta sheets in the ANKZF1-MTS.

**Figure 3:**
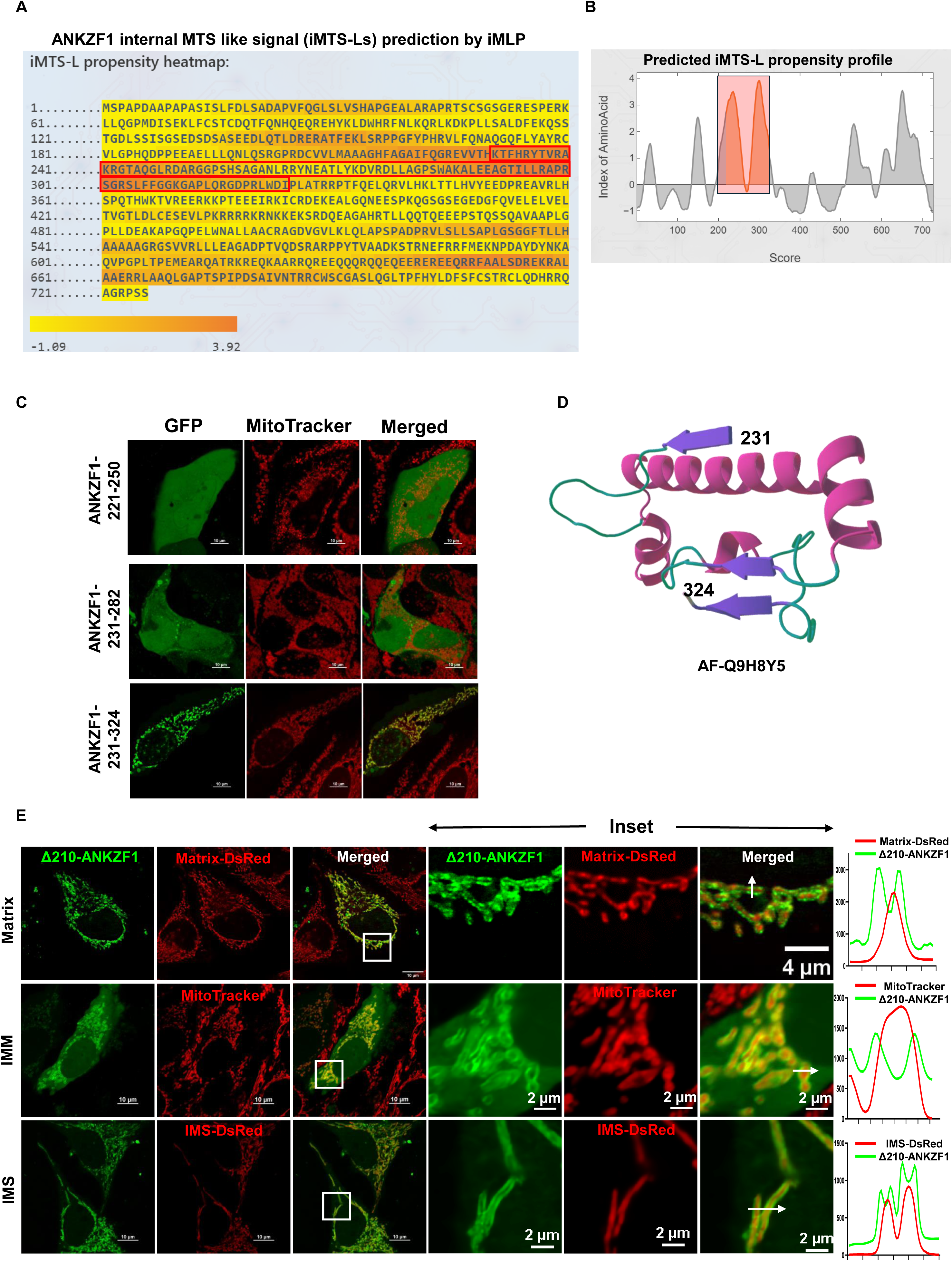
ANKZF1 N-terminal truncations localize to the outer mitochondrial membrane and Full-length ANKZF1 localizes to mitochondria occasionally. **A and B.** Predicted internal MTS-like signals (iMTS-Ls) in the ANKZF1 primary sequence by iMLP tool. In the iMLP output, the propensity of iMTS-Ls formation is shown as heatmap and suggested the presence of multiple iMTS-Ls in the ANKZF1 sequence. Two prominent iMTS-Ls lying between 231^-^325 amino acids of ANKZF1 are highlighted in a red box. **C.** Three peptide sequences of ANKZF1 (corresponding to 221-250, 231-282 and 231-324 amino acids) were fused with GFP and were expressed in HeLa cells. Images captured by confocal microscopy show 221-250-ANKZF1-GFP and 231-282-ANKZF1-GFP remain in the cytosol while 231-324--ANKZF1-GFP translocates to mitochondria and co-localizes with MitoTracker. **D.** 231-324 region of ANKZF1 was extracted from alpha fold predicted full-length structure, showing the presence of a beta-hairpin loop followed by the alpha-helical region. **E.** Δ210-ANKZF1-GFP was imaged with matrix-targeted DsRed protein, inner mitochondrial membrane-targeted MitoTracker and IMS-targeted DsRed protein. Zoomed inset shows the presence of ANKZF1 truncation mutant on the outer mitochondrial membrane. The line profile of each inset show two peaks of GFP signal while a single peak of red signal.

Interestingly, when we checked the images of mitochondria-localized Δ210-ANKZF1, we found the protein is present at the periphery of mitochondria (Figure 3E). We co-expressed matrix or IMS (Inter-membrane Space)-targeted dsRed proteins along with Δ210-ANKZF1-GFP or co-stained Δ210-ANKZF1-GFP expressing cells with Mitotracker Red which localizes to inner mitochondrial membrane. In all three conditions, we found that the green signal surrounds the red signal, and red and green signals do not overlap (Figure 3E). All the images were taken in confocal microscope and with the resolution of the confocal microscope, it is difficult to separate intra-mitochondrial structures. However, the non-overlapping signals of ANKZF1 with IMS, inner membrane or matrix-targeted probes, indicates that ANKZF1 is plausibly targeted to outer mitochondrial membrane. In summary, we show that two consecutive iMTS-Ls at residues 231-324 of ANKZF1 together act as an independent mitochondrial targeting signal and target the protein to the outer mitochondrial membrane.

### The N-terminal 72 residues of ANKZF1 prevent the mitochondrial localization of the protein and hinder the translocation of **Δ**73-ANKZF1 when expressed separately

To check the mitochondrial translocation inhibitory potential of the N-terminal segment (NTS) of ANKZF1, we cloned its first 72 amino acids at the N-terminus of mCherry (Figure 4A) and co-expressed with Δ73-ANKZF1-GFP (Figure 4E and Figure 4F). Strikingly, upon co-expression with 1-72-ANKZF1-mCherry, mitochondrial localization of Δ73-ANKZF1 was hindered (Figure 4B, 3^rd^ row from top). Importantly, mitochondrial targeting of another truncation mutant Δ210-ANKZF1 was not hindered by co-expression of 1-72-ANKZF1-mCherry (Figure 4B, bottom row) indicating that the masking of ANKZF1-MTS occurs by the interaction of specifically 61-72 residues with residues between 74-210 residues. Additionally, we showed that 1-72-ANKZF1 peptide could mask its own MTS by intramolecular as well as by inter-molecular protein-protein interaction and efficiently prevent the mitochondrial targeting of ANKZF1 (Figure 4B and C).

**Figure 4:**
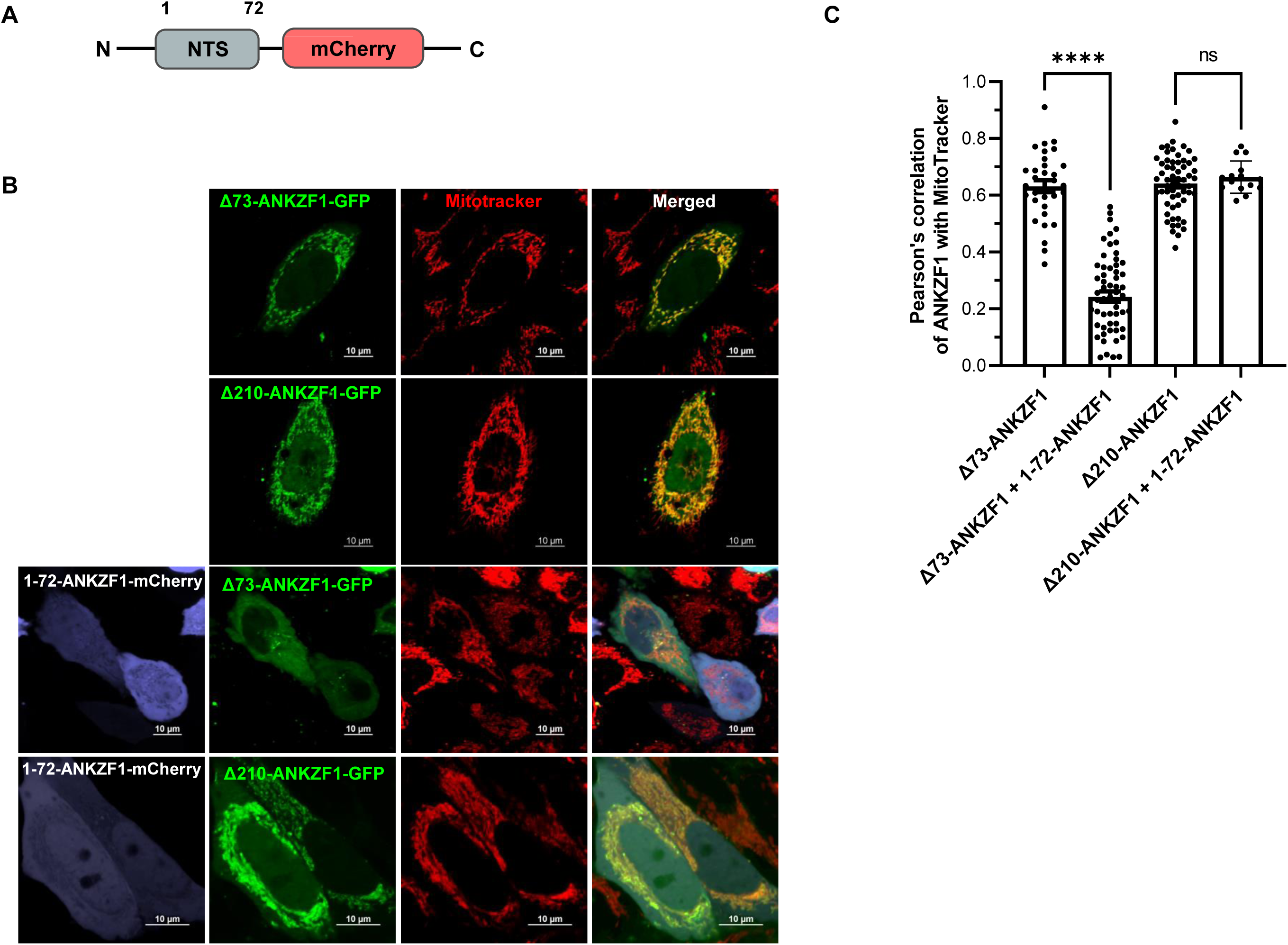
The domain-terminal 72 residues of ANKZF1 is essential for its cytosolic localization and negatively regulate the mitochondrial localization of the protein. **A.** The N-terminal 72 amino acids of ANKZF1 (N-terminal segment; NTS) were fused at the N-terminus of mCherry protein. **B.** The top two r rows show Δ73-ANKZF1 and Δ210-ANKZF1 co-localization with MitoTracker. The bottom two rows show expression of Δ73-ANKZF1 and Δ210-ANKZF1 along with 1-72-ANKZF1-mCherry. Δ73-ANKZF1-GFP shows cytosolic expression in the presence of 1-72-ANKZF1-mCherry; However Δ210-ANKZF1-GFP shows mitochondrial localization. **C.** Bar graph represents Pearson’s correlation coefficient of co-localization of ANKZF1-GFP (Δ73-ANKZF1 and Δ210-ANKZF1) and MitoTracker deep red. Values represent means ± SEM, N=3, one-way ANOVA with Bonferroni’s multiple comparisons test was performed to determine the mean differences, **** indicates P < 0.0001.

As the N-terminal 1-72 amino acids of ANKZF1 inhibited the mitochondrial localization of Δ73-ANKZF1, it was interesting to check whether ANKZF1-1-73 can generally interfere with the localization of mitochondria-targeted proteins. To answer this, we utilized three different mitochondria-targeted proteins: pAcGFP1-Mito (mitoGFP), which is targeted to the inner mitochondrial membrane; OCT-HA-GFP which is targeted to the mitochondrial matrix; and SMAC-HA-GFP, which is targeted to IMS of mitochondria (Figure S3A) (22). 1-72-ANKZF1--mCherry was co-expressed with Mito-GFP, OCT-HA-GFP, and SMAC-HA-GFP. The cells were also stained with MitoTracker Deep red and showed complete co-localization of these known mitochondria-targeted proteins with the mitochondrial network (Figure S3B and S3C) suggesting that ANKZF1-1-72 peptide does not inhibit the targeting signals of any mitochondria-targeted proteins and specifically interact and mask the MTS of ANKZF1.

As shown in the previous section, the N-terminal 72 residues of ANZKF1 block its mitochondrial localization and retain the protein in the cytosol. A similar mechanism is also followed in the case of Parkin, an E3 ubiquitin ligase and important component of the PINK-Parkin mediated mitophagy pathway. The N-terminal UBL domain of Parkin masks the Ring1 domain and inhibits the mitochondrial localization of Parkin in the absence of any stress signal eliciting mitophagy (Figure S3D) (23). We generated a ΔUBL truncation of Parkin tagged with GFP at the N-terminus (Figure S3D). The GFP-ΔUBL-Parkin shows the formation of punctate structures; however, full-length Parkin remains in the cytosol (Figure S3E). Next, we co-expressed GFP-ΔUBL-Parkin and 1-72-ANKZF1-mCherry to test if it can block the Ring1 domain similarly to UBL and inhibit the Parkin puncta formation. We found no change in the localization pattern of ΔUBL-Parkin in the presence of 1-72-ANKZF1 peptide. These results suggest that the mitochondrial translocation inhibitory capacity of 1-72-ANKZF1 peptide is highly specific for ANKZF1 and can explicitly inhibit ANKZF1 localization to mitochondria and does not block mitochondrial import of other proteins.

### Truncation of N-terminal 74 residues exposes the Mitochondria Targeting Sequence of ANKZF1

To understand the molecular basis of the negative regulation of mitochondrial translocation by the N-terminal segment of ANKZF1, an all-atom Molecular Dynamic (MD) simulation was performed for 500 nanoseconds on the AlphaFold predicted structure of ANKZF1 (UniProt ID: Q9H8Y5) (Figure 5A). The AlphaFold model suggests that the N-terminal residues are packed against the VLRF1/MTD with pLDDT scores of ∼53 and ∼41 respectively, along with a low predicted aligned error between the two regions (Figure S4). MD simulations further confirm that the N-terminal forms a stable interface with the VLRF domain, driven by hydrophobic residues that facilitate this interaction. Notably, N-terminal residues Ile14, Leu16, Phe17, Leu19, Val25, Leu29, Ser30, Leu31, Val32 are found to interact with residues Leu166, Phe167, Gln168, Asn169, Leu195, Leu196, Leu199, Arg202, Arg225, Thr 344, Leu346 with an occupancy greater than 30% (Figure 5). The distance between the interacting groups, shown in Figure S4C, remains consistently below 5 Å throughout the simulation, indicating their stability. Consistent with previous observations in the yeast ortholog, where conserved leucine residues were found to be critical, we identified three conserved leucine residues (Leu19, Leu29, Leu31) in the N-terminal interface region that are buried and likely play a pivotal role.

**Figure 5:**
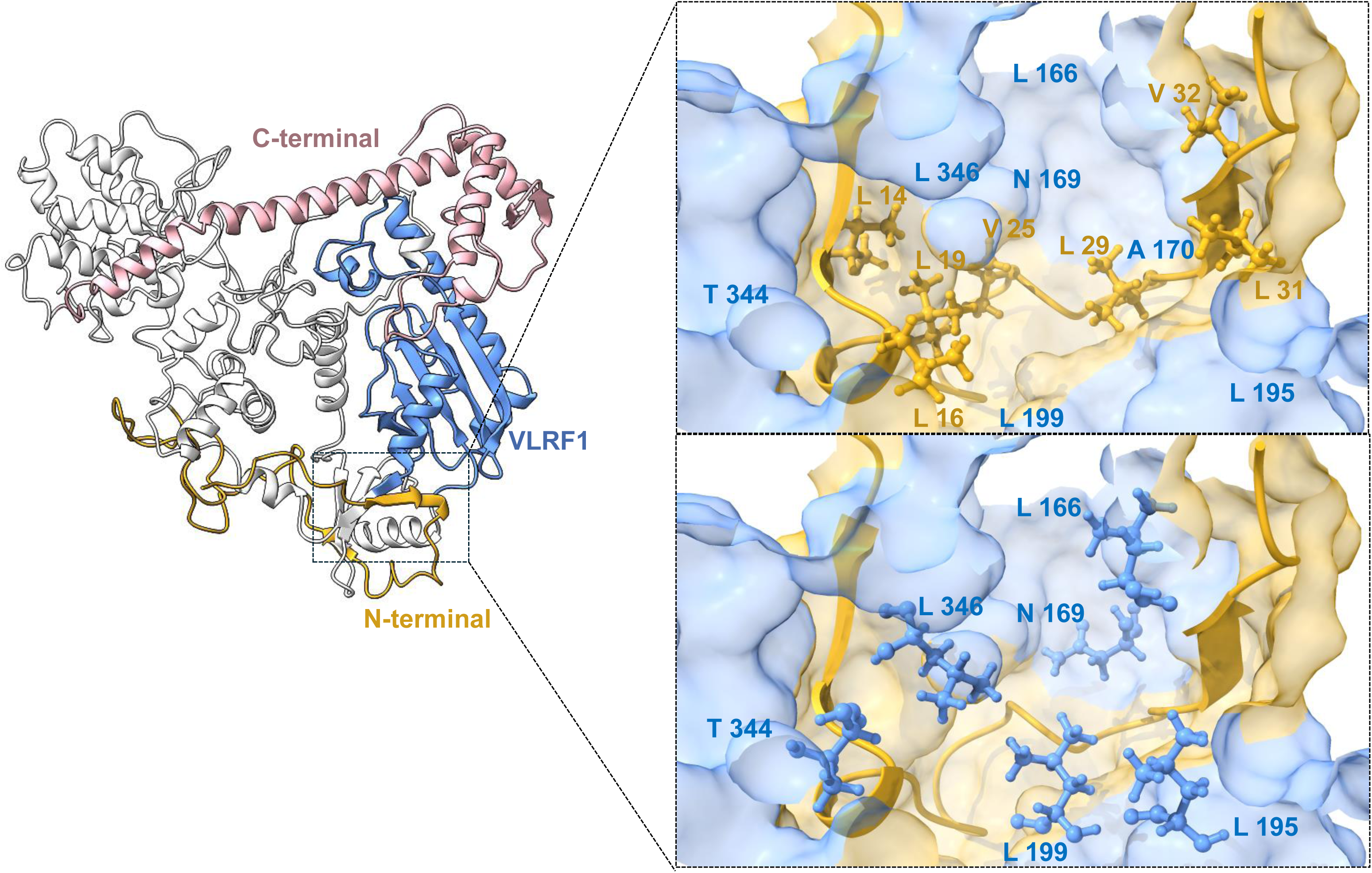
Intra-molecular interactions in ANKZF1’s N-terminal part and its VLRF domain. The left panel shows the structure of WT ANKZF1 protein after 500 ns of MD simulation. Three domains are colored differently: N-terminal (1-74, Goldenrod), VLRF1 domain (203-346, Cornflower Blue) and C-terminal domain (600-726, rosybrown). The right panel depicts the interactions between the N-terminal and VLRF1 domain. Residues Ile14, Leu16, Leu19, Val25, Leu29, Leu31, Val32 are found to interact with residues Leu166, Asn169, Leu195, Leu199, Thr 344, Leu346. For clarity, ball and stick representations of interacting N-terminal residues are depicted on the top left, and the VLRF1 motif is on the bottom left.

To understand the molecular basis of the negative regulation of mitochondrial translocation by the N-terminal part of ANKZF1, an all-atom MD simulation was performed on the AlphaFold structure of the N-terminal 74 amino acid truncated form of ANKZF1, for 500 ns. The structure obtained after 500 ns of the simulation is depicted in Figure 6A, and the corresponding root mean square deviation (RMSD) of alpha carbon atoms with respect to WT protein is shown in Figure 6B. Truncated protein was observed to display large structural deviations. Truncation of the N-terminal residues unmasks the surface of the VLRF1 domain, essential for mitochondrial translocation, and induces significant structural rearrangements. This includes disruption of interactions between the C-terminal part and the VLRF1 domain, as well as the opening of the C-terminal region (Figure 6C). Importantly, interactions in the region 203-346 amino acid residues such as Glu642-Lys241, Pro678-Arg330, Arg712-Asp274, Asp716-Arg235, which were present in the WT protein, are lost upon truncation. These structural changes lead to the exposure of the MTS, located at residues 231-240, as evidenced by solvent-accessible surface area calculations (Figure 6D), thereby facilitating the protein’s translocation to the mitochondria.

**Figure 6:**
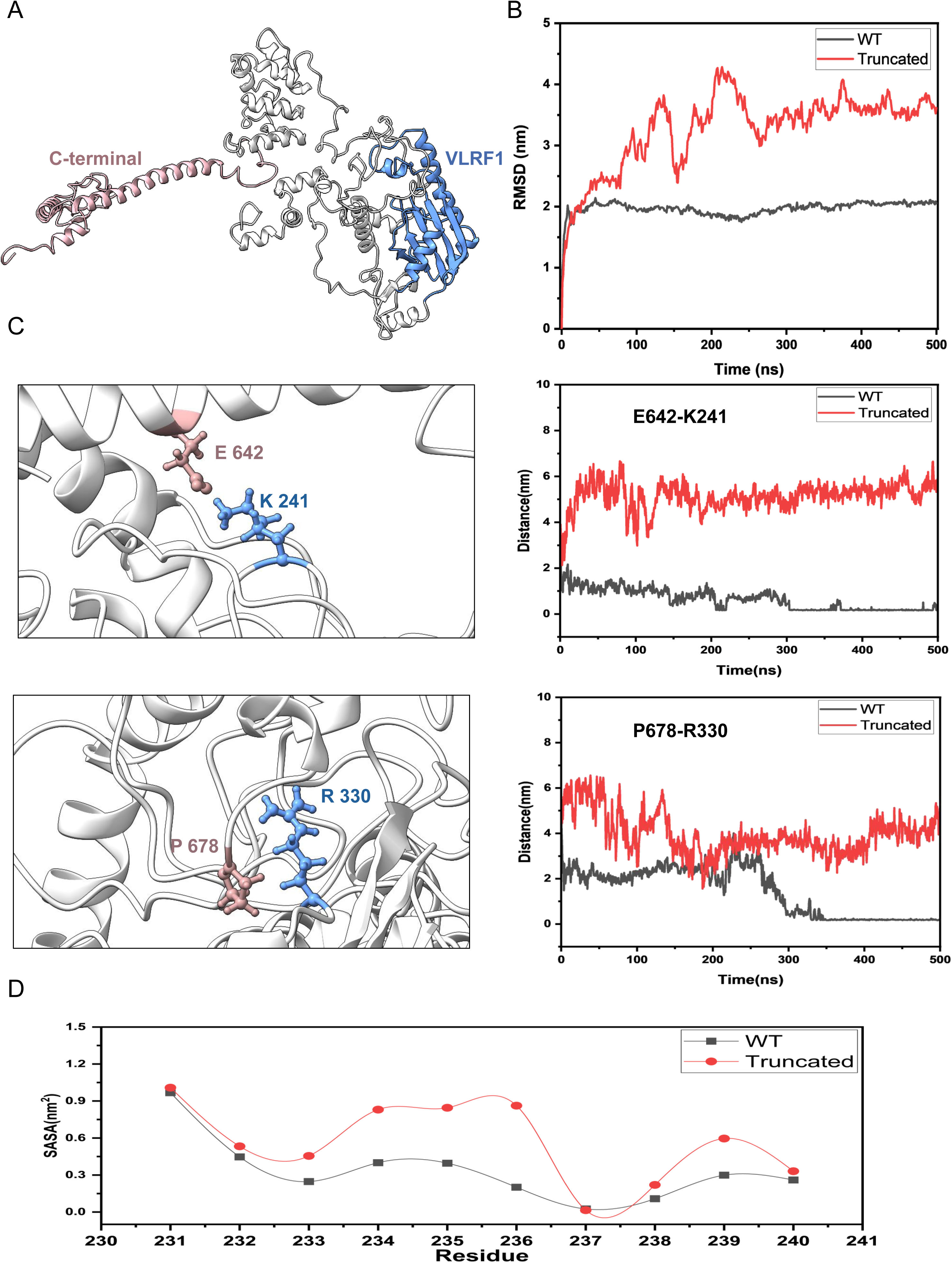
Truncation of N-terminal 74 residues exposes the internal mitochondria targeting signal of ANKZF1. **A.** Structure of N-terminal 74 residue truncated ANKZF1 protein obtained after 500 ns of MD simulations. Compared to the WT protein, the C-terminal is opened in the truncation mutant. **B** RMSD of alpha carbons for both the wild type and truncated protein over the course of a 500ns simulation. **C** 3D representation of the two key pairwise interactions between C-terminal and VLRF1 domain in the wild-type protein. These interactions are lost upon truncation of the N-terminal 74 residues. The minimum distance between the interacting pairs is shown alongside. **D** Total solvent accessible surface area of the residues from Lys231 to Ala240 for both truncated and wild-type proteins are shown.

## Discussion

In this work, we have identified and delineated the internal mitochondrial targeting signal of the human orthologue of yeast Vms1, ANKZF1. Although Vms1 was shown to localize to mitochondria during rapamycin treatment (16) or oxidative stress (14), we could not find ANKZF1 to localize to mitochondria during equivalent treatment or stress. However, we have occasionally observed ANKZF1 to localize to mitochondria (Figure S1A), although we could not identify the condition that triggers its mitochondrial localization. In this study, we have pinpointed the exact region of the protein required for its mitochondrial targeting.

### Comparison of Vms1 and ANKZF1’s internal mitochondrial targeting sequence

In a previous study, Heo et al have delineated the mitochondrial targeting domain (MTD) of yeast Vms1 (16). The authors showed that residues 182-417 of Vms1 constitute the MTD and N-terminal 182 residues inhibit the mitochondrial localization of ANKZF1. In the current study, using a series of truncation mutants, we show that an evolutionarily conserved short stretch of amino acids, 231-240 residues of ANKZF1, are indispensable for its mitochondrial localization. Furthermore, we showed that 94 residues (amino acids 231-324) constitute an independent mitochondrial targeting signal of ANKZF1. The size discrepancy of Vms1 and ANKZF1’s MTD is possibly due to the lack of shorter truncation mutants used by Heo et al. In our study, we have shown a much smaller stretch, which adopts an alpha-helical structure mainly and forms the internal mitochondrial signal sequence.

### Masking of internal mitochondrial targeting sequence used by proteins for various purposes

In this study, we have shown that the N-terminal segment of ANKZF1 interacts with the MTD and normally prevents the protein from localizing to mitochondria (summarized in Figure 7). However, due to yet unknown mechanisms, this interaction is lost, leading to conditional mitochondrial localization. In many previous studies, dually targeted proteins were reported (24). Conditional mitochondrial targeting is exhibited by many cellular proteins. Mitochondrial targeting at specific conditions is achieved by employing varied mechanisms. Li et al. figured out the dual localization pattern of APE1, which commonly resides in the nucleus; however, APE1 localizes to the mitochondria during oxidative stress. (25). Similarly, different isoforms of proteins as a result of alternate splicing events also occasionally become a deciding factor for the dual localization of some proteins. For example, Shc (Src homology and collagen) protein has three reported isoforms; the largest isoform is ∼ 66 kDa in size (p66Shc), and it shows cytosolic localization. However, a ∼46 kDa isoform, p46Shc, which lacks the N-terminal in contrast to the p66Shc, translocates to the mitochondria (26). Importantly, the *ANZKF1* gene contains 14 exons, which eventually leads to the formation of a 726 amino acid long protein, which we have termed as full-length wild-type ANKZF1. There are six different transcripts reported in the Ensembl database for *ANKZF1*. One of these transcripts results in the synthesis of full-length ANKZF1 (726 amino acids) and the second one forms a 516 amino residues long protein. Interestingly, the second protein (516 amino acids) synthesized from the second transcript starts from the 211^th^ amino acid of the wild-type ANKZF1 protein and the truncation mutant Δ210-ANKZF1 used by us corresponds to this form of the protein. However, when we checked for alternate splicing (data not shown) of *ANKZF1* gene, we could not find the formation of smaller transcripts under physiological conditions in HeLa cells. This may be due to tissue-specific or condition-specific expression of the other transcripts. In the conditions where ANKZF1 smaller transcripts are expressed, the 516 amino acid residue protein corresponding to Δ210-ANKZF1 will be translated. As shown by us, this form of the protein is exclusively localized mitochondria indicating mitochondria-specific activities of the protein.

**Figure 7:**
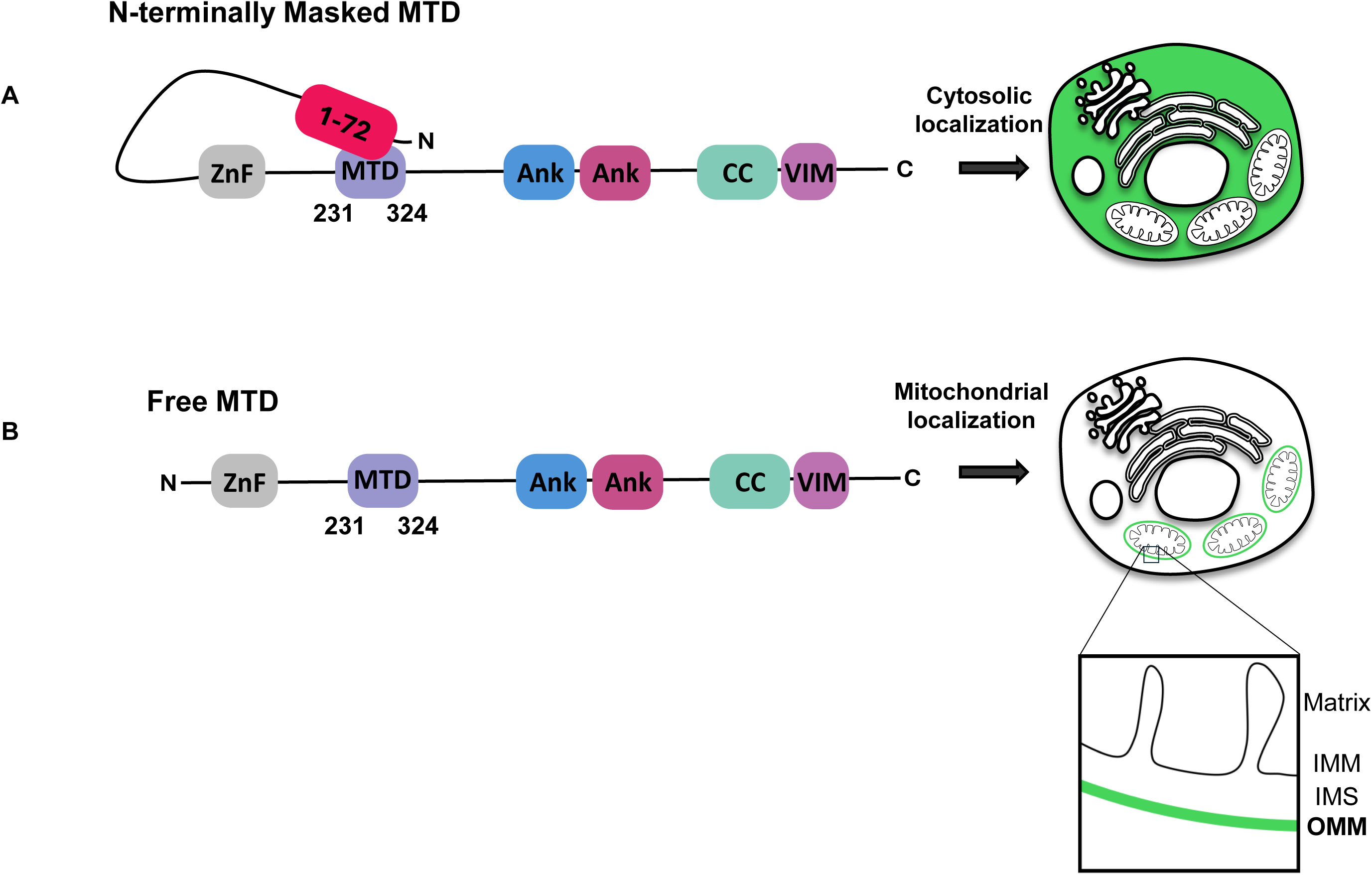
Intramolecular interaction of internal MTS of ANKZF1 with its N-terminus Inhibits its Mitochondrial Localization. **A.** Schematic diagram showing full-length WT ANKZF1 remains in the cytosol by intramolecular interaction; the N-terminal segment (1-72 amino acids) masks the internal mitochondrial targeting sequence that results in its cytosolic localization. **B.** N-terminal truncated variants carrying a free unmasked MTD resulting in its mitochondrial localization.

### Possible reasons for ANKZF1’s mitochondrial localization

As mentioned earlier, we have occasionally observed the WT ANKZF1 to localize to mitochondria although the exact cellular condition that triggers this altered intra-molecular interaction leading to exposure of internal MTS of ANKZF1 remains elusive. However, we hypothesized that certain cellular stresses like oxidative stress, as shown by its yeast ortholog, Vms1, could be one such condition. However, we did not observe mitochondrial localization of ANKZF1 by treatment with commonly used oxidative stress-imparting small molecules like H_2_O_2_ or paraquat. Even with treatment with common stressors mimicking various other environmental stresses in HeLa, HEK293T, or A5429 cells, we could not recapitulate the mitochondrial localization of wild-type ANKZF1. Recent literature has designated ANKZF1 as a transcription factor (TF), the altered expression of which, along with various other TFs, was proposed to be a predictor for colorectal cancer survival (11). Keeping this in mind, we posit that ANKZF1 may work as a transcription factor during certain stresses within mitochondria as shown for other transcription factors like Rox1 in yeast (27). These proposed reasons for ANKZF’1 conditional mitochondrial localization are only at the hypotheses stage currently and require extensive experimental validation in the future.

## Materials and Methods

### Cell Culture, Transfection and Treatments

HeLa cells were cultured in DMEM medium containing 10% FBS (Gibco) and 1% Penicillin-Streptomycin and grown at 37°C with 5% CO_2_. All plasmid DNAs were transfected in HeLa cells by Lipofectamine 2000^TM^ (Invitrogen) using the manufacturer’s standard protocol. 200nM Mitotracker deep red was used for 30 minutes to stain the mitochondria before the confocal microscopy.

### Constructs and Cloning

Plasmid constructs used in this study include the following: pCMV0-OCT-HA-eGFP (Addgene, #67479), pCMV0-SMAC-HA-eGFP (Addgene #67486), both the plasmids were a gift from Richard Kahn (28). mCherry-Parkin (Addgene #23956) was a gift from Richard Youle (29), and Mito BFP (Addgene, #55248) (30) was a gift from Michael Davidson. Δ73-ANKZF1-GFP, Δ210-ANKZF1-GFP, Δ220-ANKZF1-GFP, Δ230-ANKZF1-GFP, Δ240-ANKZF1-GFP, Δ250-ANKZF1-GFP, Δ330-ANKZF1-GFP, Δ370-ANKZF1-GFP, and Δ410-ANKZF1-GFP constructs were generated in the lab by using standard restriction digestion-based molecular cloning technique, and desired genes were digested with XhoI and KpnI restriction enzymes (NEB) and were ligated to pEGFP-N1 (Clontech) empty vector backbone. Matrix DsRed and IMS DsRed constructs were generated in the lab by fusing the Matrix and IMS targeting signals from pCMV0-OCT-HA-eGFP and pCMV0-SMAC-HA-eGFP respectively into DsRed-N1 plasmid vector (Clontech) by using XhoI and HindIII restriction digestion based cloning approach. Similarly, 1-72-ANKZF1 mCherry was generated by inserting the 1-72 amino acid coding sequence in the XhoI-KpnI sites in mCherry-N1 plasmid vector. The mito-PMD-BFP construct was previously generated in our lab (31). GFP-ΔUBL-Parkin truncated variant of Parkin was generated by cloning the corresponding gene sequence at HindIII-BamHI sites of pEGFP-N1 (Clontech) vector.

### Western Blot

Cells were lysed by RIPA lysis buffer, protein concentration was estimated and resolved in 10-12% PAGE (polyacrylamide gel electrophoresis) followed by western transfer of proteins from gel to PVDF membrane (Millipore). Membranes were blocked using 5% skimmed milk or 5% BSA dissolved in TBS-T for 1-2 hours on a slow rocker at room temperature. PVDF membranes were incubated with primary antibody diluted in 5% skimmed milk or 5% BSA (TBS-T) overnight on a slow rocker at 4°C, followed by washing and secondary antibody incubation. Lastly, blots were developed either in the Protein Simple or Bioread Chemidoc system or in x-ray films.

### Live Cell Imaging and Image Analysis

Cells were seeded on round bottom confocal dishes, and 24 hours later, cells were transfected with respective constructs and further incubated for 24 hours in the CO_2_ Incubator. All the images were recorded in Apochromat 60X or 100X 1.4 NA oil immersion-based objective lens in a Nikon A1R MP+ Ti-E confocal microscope system, under a controlled environment (temperature 37°C, 5% CO_2,_ and humidity). 405nm, 488nm, 561nm, and 640 lasers were used to excite the signals from BFP, GFP, RFP/mCherry, and MitoTracker deep red respectively. All the post-processing of the confocal data was performed in the Nikon-Essentials analysis software. Co-localization analysis and line profile assessment were also performed in the Nikon Essentials either for whole cell or desired ROI.

### MTS Prediction and Multiple Sequence Analysis

ANKZF1 Mitochondrial Targeting Sequence (MTS) were predicted by using online tool, MULocDeep (https://www.mu-loc.org/) and iMTS-Ls were predicated by iMLP online tools (https://csb-imlp.bio.rptu.de/). 211-282-ANKZF1 structure was extracted from Alphafold Predicted full length structure (AF-Q9H8Y5-F1) by using PyMOL. All the multiple sequence alignments of ANKZF1 sequences were performed in Clustal Omega and T-COFFEE multiple sequence alignment servers.

### Statistics

Initially all the datasets were assessed for their normal distribution by Anderson-Darling test and Shapiro-Wilk test. If the dataset followed the normal distribution, One-way ANOVA was performed to check the statistical significance, followed by the Bonferroni post hoc correction for the multiple comparisons. When the datasets were not following the normal distribution, a non-parametric alternative of ANOVA, the Kruskal-Wallis test, was employed, followed by Dunn’s test to analyze the pairwise comparisons between the groups. For all the analyses, statistical significance was set at p < 0.05, and all the error bars indicate mean ± SEM. All the statistical analyses were performed using GraphPad Prism.

### All-atom Molecular Dynamics Simulations

The structure of the human tRNA endonuclease ANKZF1 protein (UniProt ID: Q9H8Y5) was retrieved from the AlphaFold database (32,33). All-atom Molecular Dynamics (MD) simulations were conducted using GROMACS 2020.4 version (34). Two systems were prepared for simulation: one with the complete protein structure (WT) and another with residues 1 to 74 from the N terminal being removed (truncated), in line with our experimental setup. The CHARMM36 force field was applied for simulations, with standard protonation states assigned to ionizable residues at pH 7 using PropKa (35,36). Solvation was performed using the TIP3P water model (37) within a cubic box, maintaining a 10 Å buffer between the protein complex surface and the box edge. NaCl was added to neutralize the systems.

Energy minimization was done using the steepest descent method until the energy gradient dropped below 1000 kJ mol-1 nm-1. Equilibration was achieved in two steps: using NVT ensemble followed by NPT ensemble. The temperature was maintained at 300 K and pressure at 1 bar using V-rescale thermostat (38) and the Parrinello–Rahman barostat (39) respectively. The production run was performed in the NPT ensemble, where the Nose-Hoover thermostat was used to control the temperature and that of Parrinello-Rahman barostat for the pressure (40,41).

Coordinates were saved every 100 ps with a time step of 2 fs. Each system was simulated for 500 ns. Hydrogen bond and van-der Waals interactions between N-terminal (1-74) and VLRF1 motif (203-346) and those between VLRF1 motif (203-346) and C-terminal (600-726) were computed. Prominent interactions with an occupancy of >30% of the simulation time were analyzed. Van der Waals interactions were identified when the fraction of overlap between the van der Waals radii of two atoms was ≤0.01, while hydrogen bonds were defined by a donor-acceptor distance ≤3.9 Å, a hydrogen-acceptor distance ≤2.5 Å, and a D-H···A angle ≥120°.

## Acknowledgements

KM acknowledges the funding support from the Science and Engineering Research Board (SERB), Government of India, for Core Research Grant (SERB/CRG/2022/006517) and SNIoE core funding. MA acknowledges the SNIoE PhD fellowship and fellowship from ICMR SRF Grant (2021-14421/CMB-BMS). MA and KM acknowledge the SNU DST-FIST grant [SR/FST/LS-1/2017/59(c)] for the confocal microscopy facility. We thank Rajan Singh for his help at the confocal microscopy facility at SNIoE.

## Author Contribution

The work was conceived by KM and MA. Cell culture, molecular biology and imaging experiments were done by MA and BM. DS and NM performed the MD simulations and prepared the relevant figures. Imaging data were analyzed by MA and KM. KM supervised the work; MA and KM wrote the manuscript.

