## Supplementary figures and images for "The N-terminal Segment of the Human ANKZF1 negatively regulates its internal mitochondrial targeting signal to prevent its mitochondrial localization"

### Supplementary Figure 1

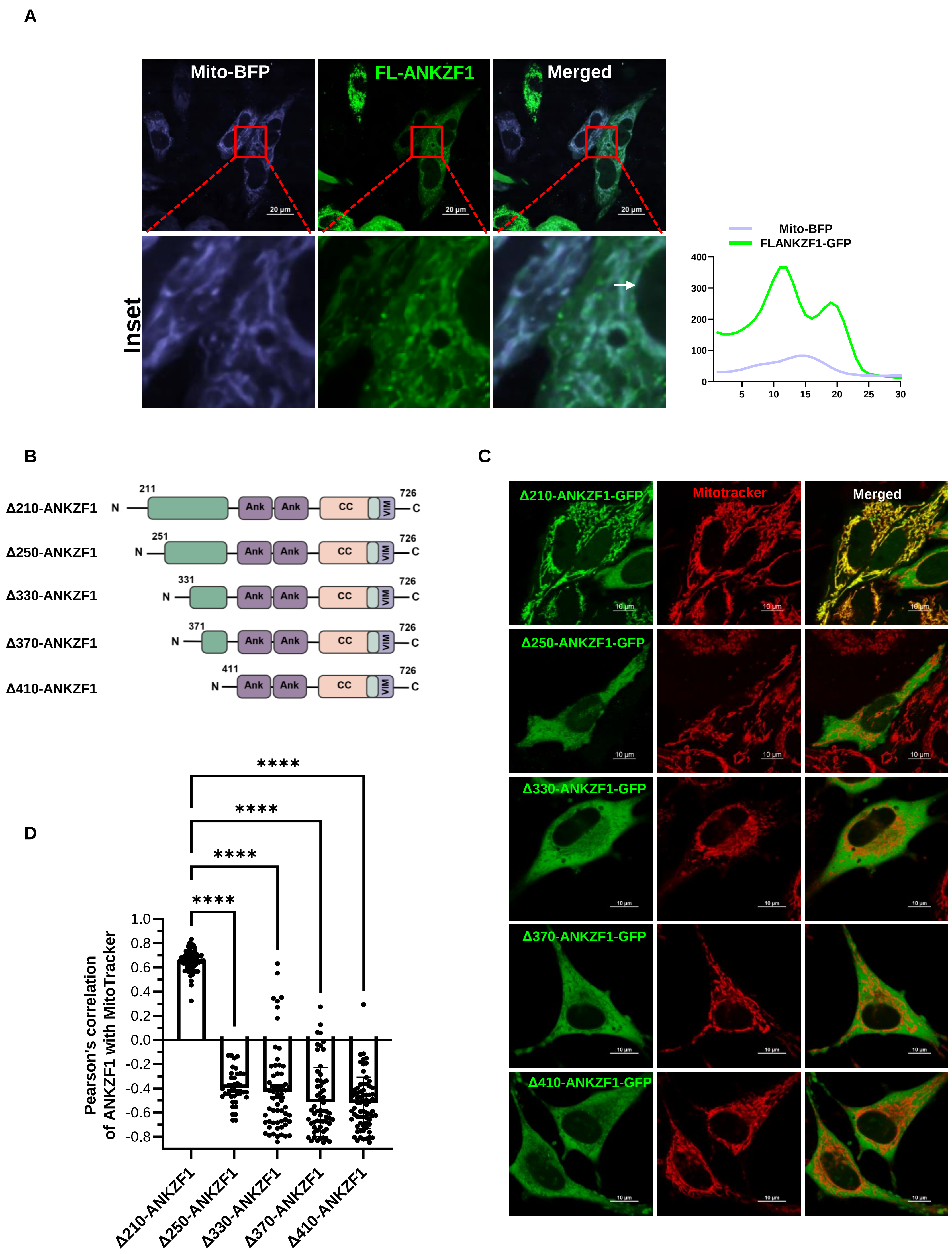

### Supplementary Figure 2

Figure S2

**A**

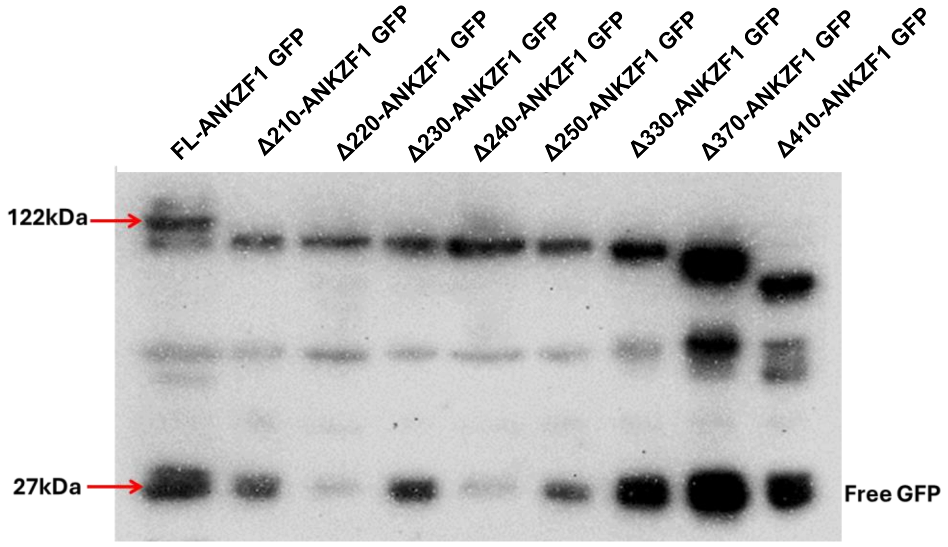


**B**

MULocDeep: ANKZF1 localization prediction at sub-cellular level

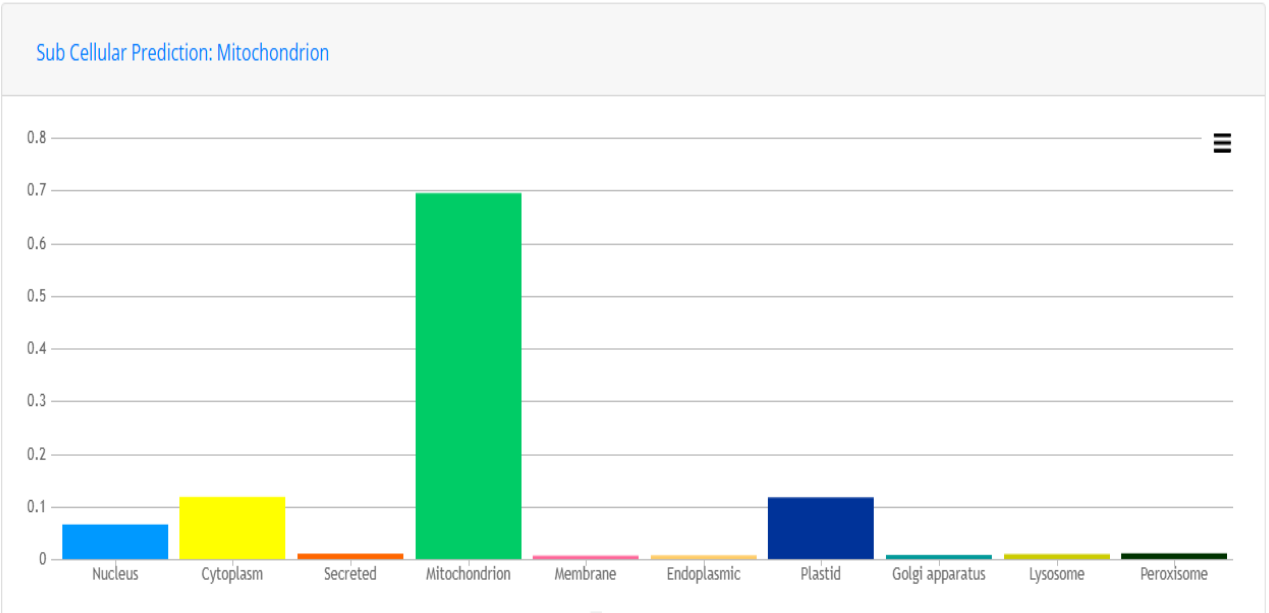

### Supplementary Figure 3

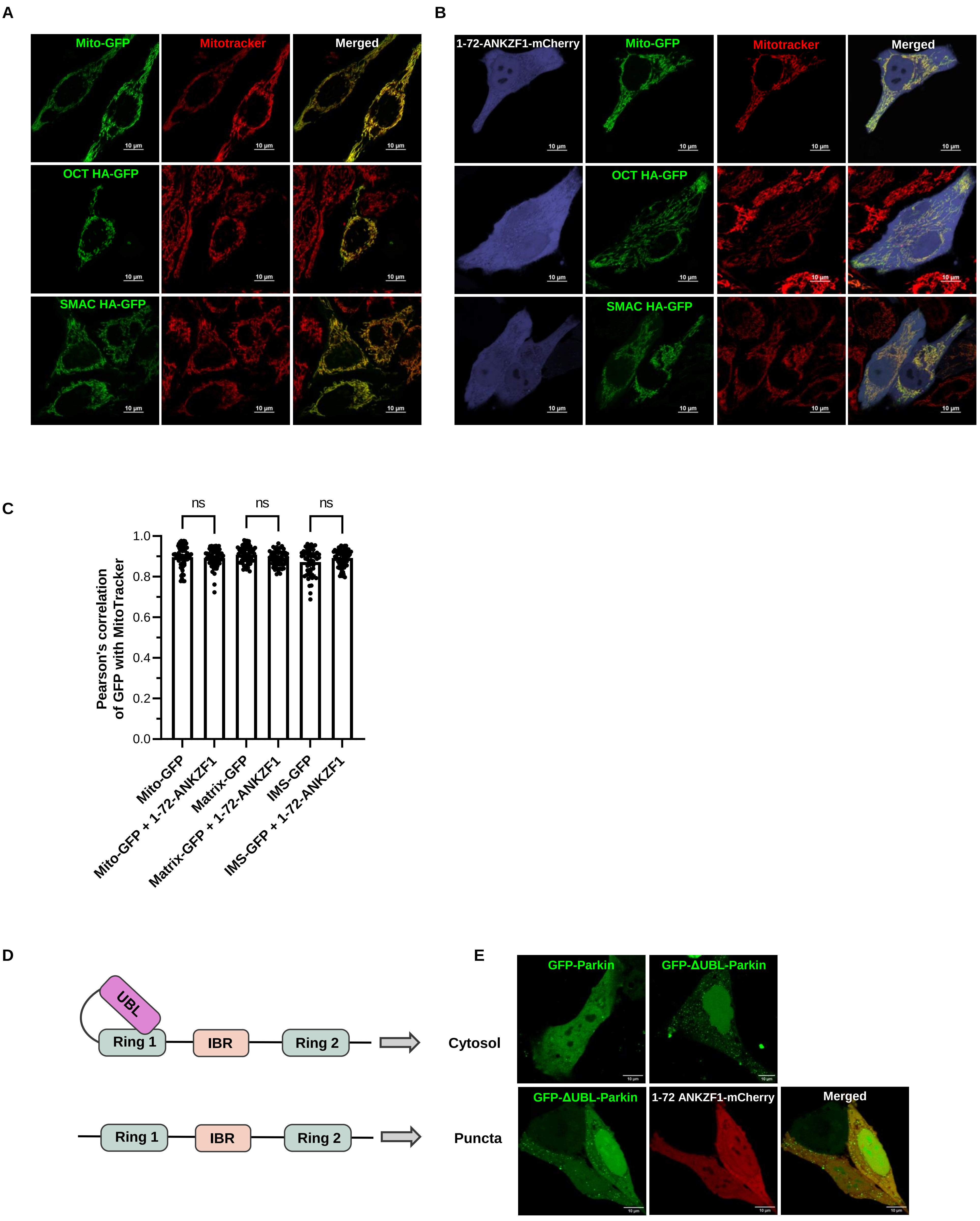

### Supplementary Figure 4

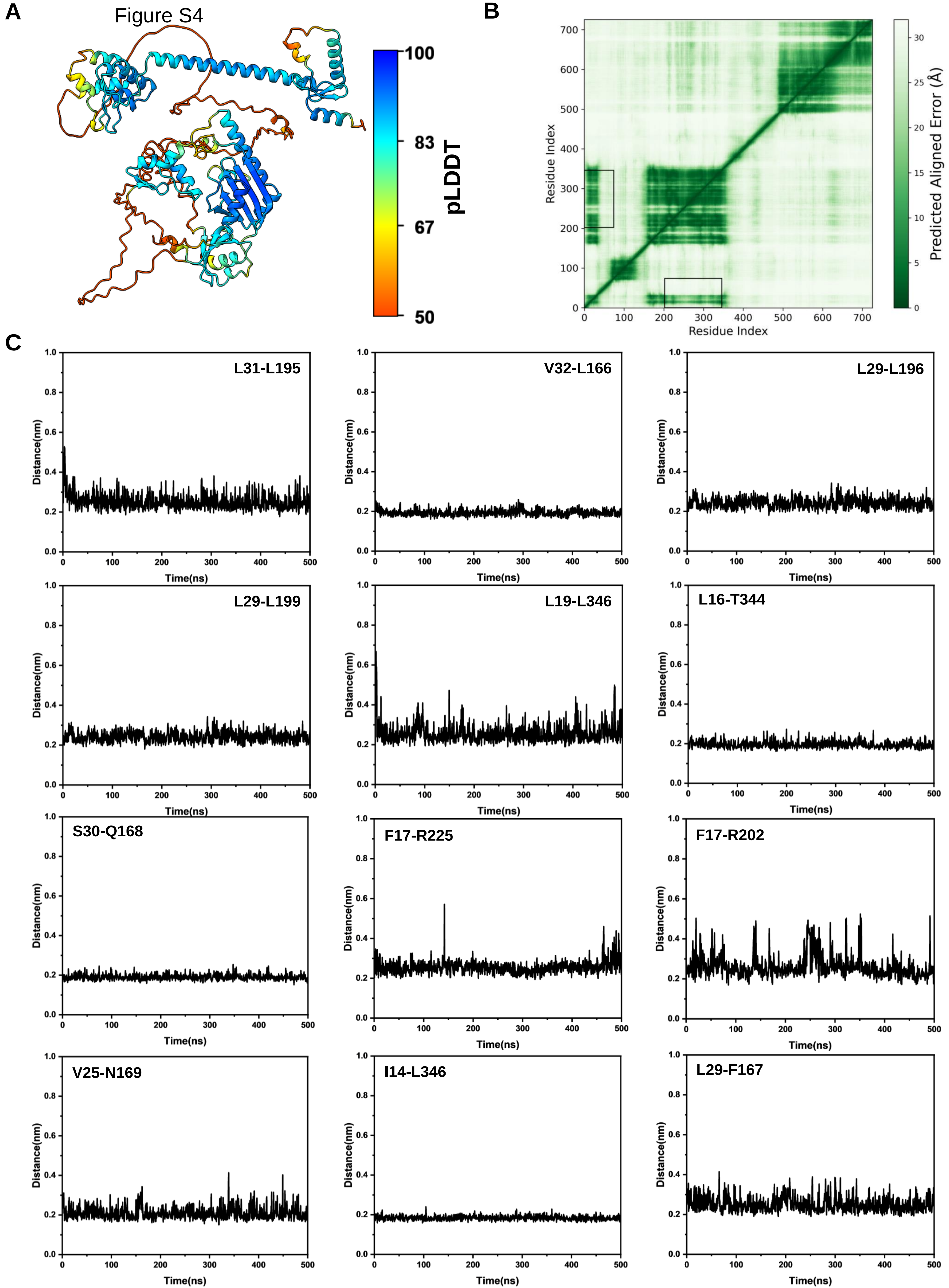
