## Supplementary Information for "The N-terminal Segment of the Human ANKZF1 negatively regulates its internal mitochondrial targeting signal to prevent its mitochondrial localization"

**Supporting Information Figure legends**

**Figure S1**

**A.** FL ANKZF1 was found to be co-localized with mito-BFP in HeLa cells occassionaly. Line profile showing the colocalized signal of ANKZF1-FL and Mito-BFP. **B.** Schematic representation showing ANKZF1 N-terminal truncations construct of Δ210-ANKZF1, Δ250-ANKZF1, Δ330-ANKZF1, Δ370-ANKZF1, and Δ410-ANKZF1 generated to determine the mitochondrial targeting domain. **C.** Δ210-ANKZF1 truncation mutant localizes to mitochondria as described in Figure 1B, here used as a positive control for mitochondrial localization. All the other truncations of ANKZF1 starting from Δ250-ANKZF1 to Δ410-ANKZF1 shows a cytosolic localization. MitoTracker deep red was used to stain the mitochondria. **D.** Bar graph showing Pearson’s correlation coefficient of co-localization of ANKZF1-GFP (Δ210-ANKZF1, Δ250-ANKZF1, Δ330-ANKZF1, Δ370 -ANKZF1, and Δ410-ANKZF1) and MitoTracker deep red. Values represent means ± SEM, N=3, Kruskal-Wallis test with Donn's multiple comparisons test was performed to determine the statistical significance.

**Figure S2**

**A.** Western blot of the WT full-length ANKZF1-GFP and all the truncated variants of ANKZF1 tagged with GFP (Δ210-ANKZF1, Δ220-ANKZF1, Δ230-ANKZF1, Δ240-ANKZF1, Δ250-ANKZF1, Δ330-ANKZF1, Δ370-ANKZF1 and Δ410-ANKZF1) showing the formation of final product at the desired molecular weight, suggesting the formation of functional proteins of expected sizes. **B.** ANKZF1 sequence from 221-324 was scanned through MULocDeep server to identify the presence of mitochondria targeting domains. It shows a ~70% probability of localizing into mitochondria.

**Figure S3**

**A.** Mito-GFP, OCT-HA-GFP (mitochondrial matrix targeted GFP), SMAC-HA-GFP (mitochondrial IMS targeted GFP) were expressed in HeLa cells and the cells were stained with MitoTracker deep red. Merged channel shows the co-localization of GFP signals with MitoTracker signal suggesting the localization of these proteins into the mitochondria. **B.** Mito-GFP, OCT-HA-GFP, and SMAC-HA-GFP were co-expressed with 1-72-ANKZF1. MitoTracker staining was done before imaging, and all the panels showed GFP signal co-localized with MitoTracker signals, suggesting 1-72-ANKZF1 does not inhibit the localization of any other mitochondria targeted proteins. **C.** Bar plot shows the co-localization status of GFP and MitoTracker in the presence of 1-72-ANKZF1. Values represent means ± SEM, N=3, and one-way ANOVA with Bonferroni's multiple comparisons test was performed to determine the mean differences. **D.** schematic representation of the domain map of full-length Parkin protein and ΔUBL-Parkin. **E.** Upper panel shows the expression pattern of full-length Parkin showing its cytosolic expression. However, ΔUBL-Parkin shows a punctate pattern. Bottom panel, ΔUBL-Parkin co-expressed with 1-72-ANKZF1, but no change in the localization pattern was observed.

**Figure S4**

**A**. AlphaFold-predicted structure of ANKZF1 protein, colored based on pLDDT scores. **B** Predicted Aligned Error (PAE) plot showing high confidence structural prediction between the N-terminal 1-74 residues with VLRF1 domain (203-346). **C** The minimum distance between the interacting pair groups of N-terminal residues and the residues of the VLRF1 domain

**Supplementary References**
