## Supplementary Table 1 for "The N-terminal Segment of the Human ANKZF1 negatively regulates its internal mitochondrial targeting signal to prevent its mitochondrial localization"

**Supporting Information table 1.**

**List of plasmid constructs used in this paper.**

| **Construct Name** | **Vector Backbone** | **Source** | **Addgene Number** |
| --- | --- | --- | --- |
| EBFP2-Mito-7 | EBFP2 | Addgene | 55248 (1) |
| mCherry-Parkin | mCherry-C1 | Addgene | 23956 (2) |
| OCT HA-GFP | pcDNA3.1 | Addgene | 67479 (3) |
| SMAC HA-GFP | pcDNA3.1 | Addgene | 67486 (3) |
| RFP-LC3 |  | Received as a gift from Professor Oishee Chakrabarti’s lab, SINP, Kolkata |  |
| Matrix-DsRed | DsRedN1 | Generated in the lab |  |
| IMS-DsRed | DsRedN1 | Generated in the lab |  |
| Mito-PMD-BFP | EBFP | Generated in the lab |  |
| ANKZF1-FL-GFP | pcDNA3.1 | Generated in the lab |  |
| 1-72-ANKZF1-mCherry | mCherry N1 | Generated in the lab |  |
| Δ73-ANKZF1-GFP | EGFPN1 | Generated in the lab |  |
| Δ210-ANKZF1-GFP | EGFPN1 | Generated in the lab |  |
| Δ220-ANKZF1-GFP | EGFPN1 | Generated in the lab |  |
| Δ230-ANKZF1-GFP | EGFPN1 | Generated in the lab |  |
| Δ240-ANKZF1-GFP | EGFPN1 | Generated in the lab |  |
| Δ250-ANKZF1-GFP | EGFPN1 | Generated in the lab |  |
| Δ330-ANKZF1-GFP | EGFPN1 | Generated in the lab |  |
| Δ370-ANKZF1-GFP | EGFPN1 | Generated in the lab |  |
| Δ410-ANKZF1-GFP | EGFPN1 | Generated in the lab |  |
| GFP-ΔUBL-Parkin | EGFPN1 | Generated in the lab |  |
