## Supplementary Table 2 for "The N-terminal Segment of the Human ANKZF1 negatively regulates its internal mitochondrial targeting signal to prevent its mitochondrial localization"

**Supporting Information Table 2.**

**List of reagents and antibodies used in this paper.**

| **Reagents and Antibody** | **Made** | **Cat number** |
| --- | --- | --- |
| Lipofectamine^TM^ 2000 | Thermo | 11668027 |
| MitoTracker Far Red | Invitrogen | M22426 |
| Anti-ANKZF1 | Sigma | HPA035208 |
| Anti-GFP | Abcam and polyclonal antibody generated in-house | Ab183735 |
